# The miR-144/Hmgn2 regulatory axis orchestrates chromatin organization during erythropoiesis

**DOI:** 10.1101/2023.07.18.549576

**Authors:** Dmitry A. Kretov, Leighton Folkes, Alexandra Mora-Martin, Noreen Syedah, Isha A. Walawalkar, Kim Vanyustel, Simon Moxon, George J. Murphy, Daniel Cifuentes

**Affiliations:** Department of Biochemistry and Cell Biology, Boston University Chobanian & Avedisian School of Medicine, Boston, MA; School of Biological Sciences, University of East Anglia, Norwich, UK; Center for Regenerative Medicine, Boston University Chobanian & Avedisian School of Medicine, Boston, MA; Section of Hematology and Oncology, Department of Medicine, Boston Medical Center, Boston, Massachusetts, USA; Amyloidosis Center, Boston University Chobanian and Avedisian School of Medicine, Boston, Massachusetts, USA; Department of Virology, Immunology and Microbiology, Boston University Chobanian & Avedisian School of Medicine, Boston, MA

**Keywords:** microRNAs, miR-144, Hmgn2, erythropoiesis, chromatin, zebrafish

## Abstract

**SUMMARY:** Differentiation of stem and progenitor cells is a highly regulated process that involves the coordinated action of multiple layers of regulation. Here we show how the post-transcriptional regulatory layer instructs the level of chromatin regulation via miR-144 and its targets to orchestrate chromatin condensation during erythropoiesis. The loss of miR-144 leads to impaired chromatin condensation during erythrocyte maturation. Among the several targets of miR-144 that influence chromatin organization, the miR-144-dependent regulation of Hmgn2 is conserved from fish to humans. Our genetic probing of the miR-144/Hmgn2 regulatory axis established that intact miR-144 target sites in the Hmgn2 3’UTR are necessary for the proper maturation of erythrocytes in both zebrafish and human iPSC-derived erythroid cells while loss of Hmgn2 rescues in part the miR-144 null phenotype. Altogether, our results uncover miR-144 and its target Hmgn2 as the backbone of the genetic regulatory circuit that controls the terminal differentiation of erythrocytes in vertebrates.

## INTRODUCTION

MicroRNAs (miRNAs) are a family of small non-coding RNAs that regulate gene expression by destabilizing and repressing the translation of their target mRNAs^1^. As critical post-transcriptional regulators, miRNAs play a central role in cell differentiation and embryo development by restricting cell fate choices and regulating developmental timing^2,3^. miRNAs enact this post-transcriptional control by targeting hundreds of mRNA each^4^. However, analysis of miRNA loss-of-function mutants reveals that the bulk of expression changes come from the additional dysregulation of non-target mRNAs^1–3^. Uncovering the molecular mechanisms governing how miRNAs expand their regulatory role to additional non-target mRNAs will help to establish a more comprehensive framework of miRNA-mediated regulation of cell differentiation and development. Here we uncover and genetically probe how the post-transcriptional regulation elicited by miR-144 instructs chromatin organization during erythropoiesis by ultimately increasing the number of transcripts affected directly or indirectly by miR-144 activity.

Erythropoiesis is a highly orchestrated and dynamic process in which multipotent hematopoietic stem and progenitor cells (HSPCs) progressively differentiate to become mature erythrocytes^5^. Two vertebrate-specific miRNAs, miR-144 and miR-451, regulate erythrocyte terminal differentiation. These miRNAs are expressed from a single primary miRNA precursor whose expression is activated by the transcription factor GATA1^6^. miR-451, which accounts for up to 60%^7^ of the miRNA content in mature erythrocytes, is the only known miRNA whose processing is independent of Dicer but instead relies on the slicer activity Ago2^8–10^. miR-451 function is required for proper erythroid maturation^11,12^ and mounting the oxidative stress response^13,14^. miR-144 is involved in the regulation of globin synthesis and oxidative protection of the cell^15,16^. Our recent work also demonstrated that miR-144 establishes a negative-feedback loop with Dicer that induces the global downregulation of canonical miRNAs while promoting the Dicer-independent processing of miR-451 during erythropoiesis^17^.

Committed erythroid progenitors undergo a global repression of gene expression with the exception of erythroid-specific genes such as globins, proteins involved in membrane organization, and specific non-coding RNAs^18,19^. In all vertebrates, terminal differentiation of erythrocytes is accompanied by progressive chromatin condensation, which in mammals culminates with the extrusion of the nucleus^18,20,21^. These rearrangements in nuclear organization are needed to restrict cellular fate to the erythroid lineage and direct cellular machinery towards the synthesis of proteins necessary for the proper function of erythrocytes^22,23^. Perturbations of erythrocyte maturation can lead to anemia and myelodysplastic syndromes^24,25^.

In our current work, we conducted a phenotype-informed search to uncover miR-144 targets that orchestrate nuclear condensation during terminal erythropoiesis. We identified a gene regulatory axis, comprising miR-144 and its direct target Hmgn2, involved in chromatin organization. Disruption of the miR-144-mediated regulation of Hmgn2 recapitulates in part the impaired erythropoiesis phenotype of miR-144 mutants. Conversely, the reduction of Hmgn2 activity restores normal erythrocyte maturation in miR-144 mutants. Overall, we show how microRNAs can expand their targeting network and amplify their regulatory potential via targeting a master regulator of chromatin organization.

## RESULTS

### Loss of miR-144 leads to defects in nuclear condensation and genome-wide dysregulation of gene expression in erythroblasts

During terminal differentiation, erythroblasts progressively condense their chromatin, which culminates in the extrusion of the nucleus in mammals, or a reduction of the nuclear volume in all the other vertebrates including zebrafish^19^. Our previous morphological analysis revealed that erythrocytes isolated from zebrafish mutants with a deletion in the miR-144 locus (miR-144^Δ/Δ^) display enlarged nuclei with pronounced granular staining as compared to wild-type erythrocytes^17^, a hallmark of impaired erythrocyte maturation. To determine when miR-144^Δ/Δ^ erythrocytes first manifest nuclear defects, we conducted a time course analysis of the maturation of wild-type and miR-144^Δ/Δ^ erythrocytes. The May-Grünwald-Giemsa (MGG) staining of erythroblasts isolated from embryos at different developmental stages (30-hours post-fertilization (hpf) and 2- and 3-days post-fertilization (dpf)) shows that the morphology of miR-144^Δ/Δ^ erythroblasts is indistinguishable from wild-type siblings until 2-dpf (Figure 1A). At 3-dpf, miR-144^Δ/Δ^ erythroblasts display an increased Nucleus-to-Cytoplasm area ratio (N:C) (Figure 1A and B), typical of immature erythrocytes^12^. These results are consistent with the increased nuclear area that we found previously in erythrocytes isolated from adult fish^17^ albeit at 3-dpf peripheral blood is a mix of cells produced during primitive and definitive waves of erythropoiesis. Overall, these results indicate that the deletion of miR-144 impairs erythrocyte maturation starting at 2-dpf and persisting into adulthood.

**Figure 1.**
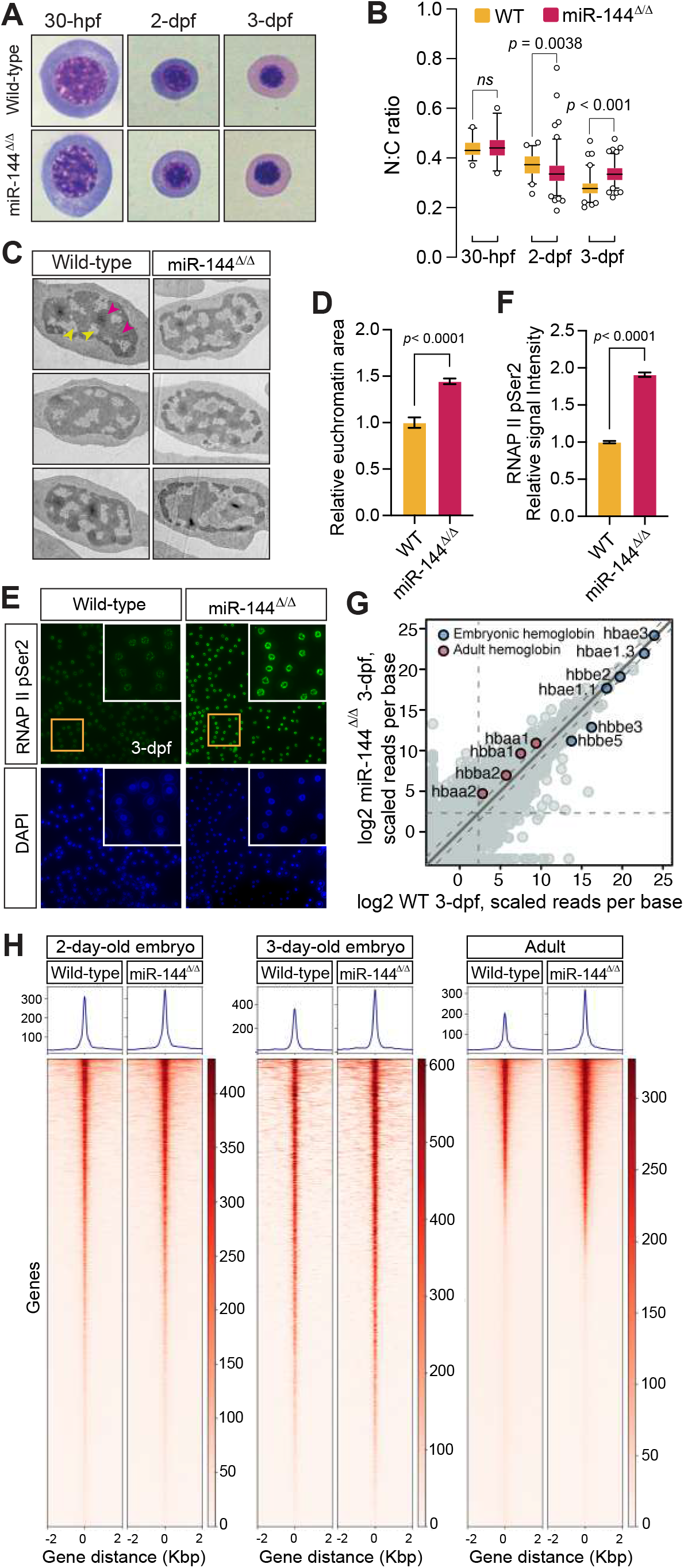
Loss of miR-144 impairs chromatin condensation during erythropoiesis. **(A)** May-Grunwald-Giemsa staining of peripheral blood cells isolated from miR-144^Δ/Δ^ and wild-type siblings at 30-hpf, 2-dpf and 3-dpf. **(B)** Quantitative analysis of Nucleus-to-Cytoplasm area ratio of erythrocytes stained in (A). Between 26 and 113 cells are analyzed in each case. *p*-values from ordinary one-way ANOVA test. Boxes enclose 5 to 95 percentiles. **(C)** Transmission Electron Microscopy of erythrocytes isolated from 3-dpf embryos (yellow arrows indicate euchromatin and magenta indicate heterochromatin). **(D)** Quantification of euchromatic regions of the nuclei from (C). Between 29 and 31 cells are analyzed in each case. *p*-values from unpaired *t-*test. Error bars represent standard error of the mean. **(E)** Immunofluorescent staining of erythrocytes isolated from 3-dpf embryos with anti-RNAP II Ser2 antibodies. **(F)** Quantification of the nuclear signal of RNAP II Ser2 from (E). 100 cells are analyzed in each case. *p*-values from unpaired *t-*test. Error bars represent standard error of the mean. **(H)** Heat-maps of ATAC-seq analysis of erythrocytes isolated from peripheral blood 2-dpf, 3-dpf and adult miR-144^Δ/Δ^ fish and wild-type siblings. 2-dpf and adult samples are analyzed in triplicates, and 3-dpf samples in duplicates. **(G)** RNA-Sequencing of erythrocytes isolated from 3-dpf. Average of 2 biological replicates is plotted. Expression of adult and embryonic *globins* is shown.

To interrogate nuclear organization in more detail, we analyzed miR-144^Δ/Δ^ erythrocytes from 2- and 3-dpf embryos by transmission electron microscopy (TEM). miR-144^Δ/Δ^ erythroblasts displayed a 1.44-fold increase of light nuclear areas (euchromatin) compared to wild-type cells only at 3-dpf (Figure 1C and D), but not at 2-dpf (Supplementary figure S1A and S1B) suggesting an impairment in heterochromatin formation during differentiation of miR-144^Δ/Δ^ cells. Following these results, we reasoned that the prevalence of euchromatin regions could be conducive to enhanced transcription^26^. To probe this hypothesis, we quantified the levels of RNA polymerase II (RNAP II) phosphorylated at Ser2 in its C-terminal domain, which is a proxy for the fraction of RNAP II engaged in transcription elongation. In concordance with our hypothesis, we observed a ∼2-fold increase in the RNAP II pSer2 signal in miR-144^Δ/Δ^ erythroblasts isolated from 3-dpf embryos as compared to their wild-type siblings (Figure 1E and F). Altogether, these results suggest that loss of miR-144 impairs chromatin condensation of erythrocytes and leads to increased transcriptional activity.

To determine if the defect in chromatin remodeling of miR-144^Δ/Δ^ erythroblasts is restricted to a few specific genomic loci or a genome-wide effect, we performed ATAC-seq analysis. We analyzed peripheral blood cells isolated from 2- and 3-dpf embryos and adult zebrafish. Analysis of ATAC-seq peaks indicated that there was no significant difference in peak accessibility between wild-type and miR-144^Δ/Δ^ erythrocytes isolated from 2-dpf animals (Figure 1H). However, we observed an increase in accessible chromatin regions in miR-144^Δ/Δ^ erythroblasts at 3-dpf and in erythrocytes of adult fish (Figure 1H) from their wild-type counterparts. This finding is in agreement with the onset of nuclear defects observed at 3-dpf (Figure 1A) and demonstrates that the loss of miR-144 has a direct effect on chromatin organization.

To determine whether the increase in open chromatin regions leads to a global increase in gene transcription, as hinted by the RNAP II pSer2 staining, we analyzed gene expression using bulk mRNA sequencing at 3-dpf (Figure 1G). Our analysis shows that the 21,013 genes with more than 5 scaled reads per base in any of the genotypes, 3,655 genes are stabilized and 1,740 genes that are destabilized more than 2-fold in miR-144^Δ/Δ^ compared to wild-type erythrocytes confirming a global trend of the increase in gene expression caused by the deletion of miR-144 (Supplementary Figure S1C).

Among the 3,655 genes expressed at higher levels in the miR-144^Δ/Δ^, only 457 contain at least one predicted miR-144 target site in their 3’UTR according to TargetScan v6.2. In addition, we observed that 272 genes of those stabilized in miR-144^Δ/Δ^ have a more than 1-fold increase in chromatin accessibility as revealed by ATAC-seq at 3-dpf. Only 38 genes with upregulated chromatin accessibility are also miR-144 targets indicating that these two regulatory programs operate largely independently. For example, we detected the expression of adult globin genes (hbaa1 and hbba1), that are not miR-144 targets, along with embryonic globins (Figure 1G). Since adult globins are only expressed in wild-type zebrafish larvae older than ∼30-days^27^, these results suggest that the increase in mRNA expression in the miR-144^Δ/Δ^ mutants is not only due to disruption of miRNA-mediated silencing and subsequent stabilization of miRNA targets but also represents a major disruption of the transcriptional program caused by alterations at the chromatin level. Moreover, in erythrocytes isolated from adult fish, we observed an even stronger correlation between an increase in chromatin accessibility and elevated gene expression in miR-144^Δ/Δ^ (Supplementary Figure S1D and S1E). This correlation coincides with the timing of the onset of chromatin condensation and highlight its important role in shaping gene expression during erythropoiesis. Moreover, these results show that the impact of miRNA-mediated regulation on gene expression goes beyond the immediate regulation of their direct targets.

### miR-144 targets several chromatin factors including Hmgn2

We hypothesized that the impaired nuclear condensation phenotype of miR-144^Δ/Δ^ erythrocytes may be driven at least in part by the gain of function of one or more of the miR-144 mRNA targets that are stabilized in the mutant fish. Our previous work demonstrated that miR-144 targets Dicer in erythrocytes, affecting the processing of miR-451 and all canonical miRNAs^17^. To test if Dicer stabilization and accompanied dysregulation of miRNA metabolism might contribute to the miR-144^Δ/Δ^ phenotype, we injected mRNA encoding Dicer in zebrafish embryos at the 1-cell stage (Figure 2B). At 2-dpf, we isolated peripheral circulating blood cells and stained the blood smears and performed MGG staining. The N:C ratio of the Dicer-injected embryos was not significantly different from the non-injected siblings (Figure 2C). These results suggest that Dicer is not directly responsible for the nuclear condensation process other than via its central role in miR-144 processing.

**Figure 2.**
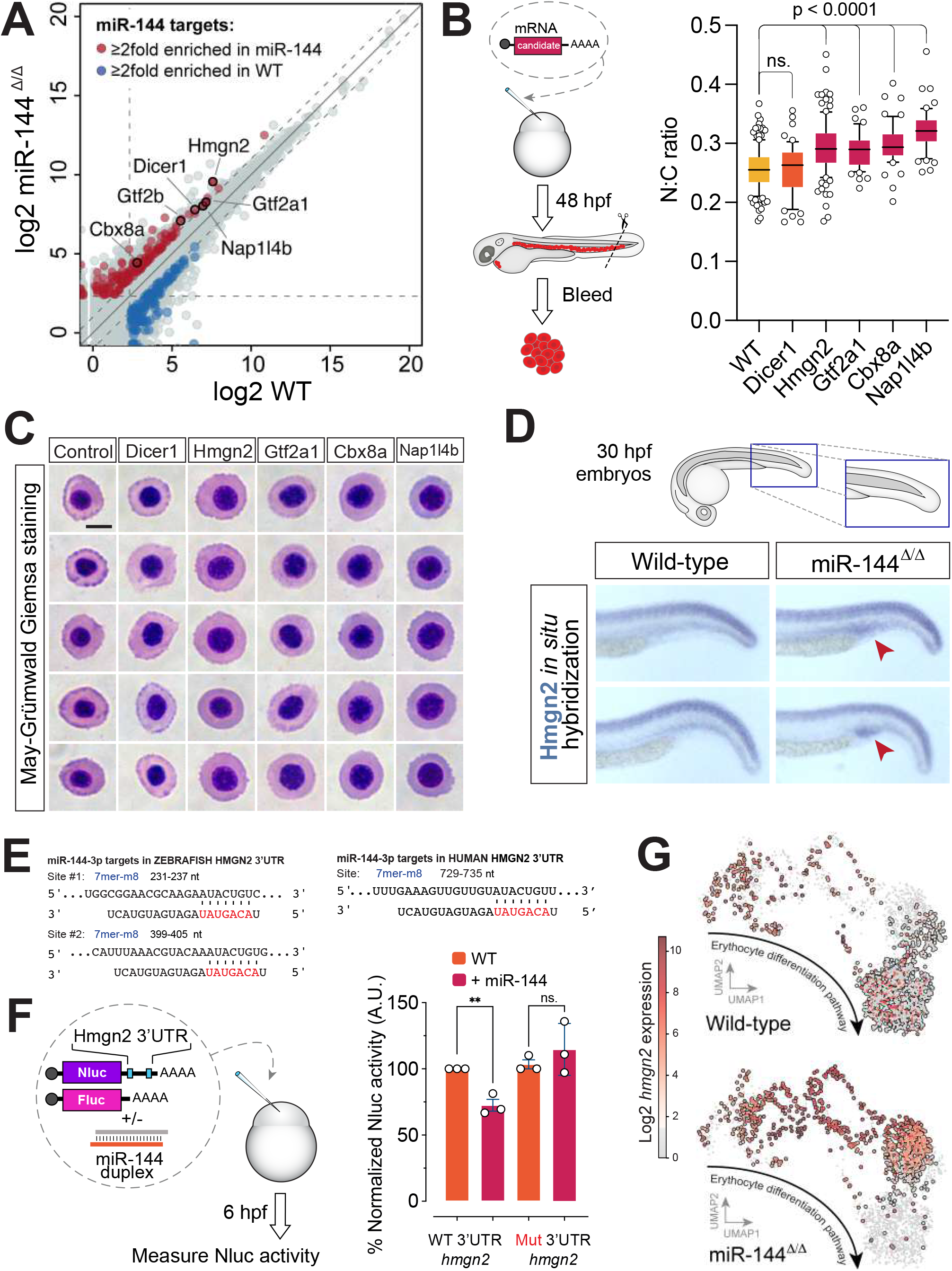
Hmgn2 is a *bona fide* miR-144 target whose expression is down-regulated in mature erythrocytes. **(A)** QuantSeq of erythrocytes isolated from 2-dpf embryos. Average of 3 biological replicates is plotted. Up-regulated (red) and down-regulated (blue) miR-144 targets are show. Genes involved in chromatin organization and transcription are outlined and labeled. **(B)** Experimental design of testing of candidate genes identified in (A) and quantification of their effect on the morphology of erythrocytes as measured by the N:C ratio after May-Grunwald-Giemsa staining (C). Between 53 and 145 cells are analyzed in each case. *p*-values from ordinary one-way ANOVA test. Boxes enclose 10 to 90 percentiles. **(C)** May-Grunwald-Giemsa staining of peripheral blood cells isolated from wild-type 2-dpf embryos over-expressing different factors. **(D)** Whole-mount *in situ* hybridization of *Hmgn2* mRNA at 30-hpf of miR-144^Δ/Δ^ fish and wild-type siblings. **(E)** Predicted miR-144-3p-v1 (according to TargetScan target sites in *Hmgn2* 3’UTR of *Danio rerio* and *Homo Sapiens*. miR-144 seed region is indicated in red. **(F)** Nanoluciferase reporter assays to validate miR-144 targeting *Hmgn2* 3’UTR. The reporter mRNA contains the Nanoluciferase (Nluc) open reading frame followed by *Danio rerio Hmgn2* 3’UTR wild-type sequence or with deleted miR-144 sites. Data represent mean ± standard deviation of three technical replicates. *p*-values from unpaired *t-*test. **(G)** Single-cell sequencing of adult zebrafish pronephros of miR-144^Δ/Δ^ and wild-type zebrafish. Only erythroid branch is shown. Expression of Hmgn2 is indicated.

To find additional candidates, we analyzed changes in gene expression in miR-144^Δ/Δ^ cells at 2-dpf because we reason that the morphological changes observed in 3-dpf embryo should be triggered by molecular alterations at earlier stages of development. We performed a gene ontology (GO) analysis on the group of stabilized miR-144 target genes (more than 2-fold upregulated and more 5 scaled reads per base (Figure 2A). Among the targets enriched for GO terms related to chromatin, we identified chromatin regulators (cdx8, napl1l4b, and hmgn2) and transcription factors (gtf2a1 and gtf2b).

To probe which of these candidates when overexpressed can recapitulate the impaired nuclear condensation of miR-144^Δ/Δ^ erythroblasts, we injected mRNAs encoding each of the candidate genes into 1-cell stage wild-type embryos. Over-expression of all new candidates increased the N:C ratio of the erythroblasts (Figure 2B and 2C). These results suggest that all these candidates participate to some degree in nuclear condensation, and that the defects observed in miR-144^Δ/Δ^ are the result of the compound gain-of-function effect of multiple genes involved in chromatin regulation. Among these candidates, Hmgn2 stands out because it: i) has the highest expression level in erythrocytes (Figure 2A), ii) has a higher N:C ratio when over-expressed (Figure 2C), iii) is a conserved miR-144 target from fish to human (Figure 2E and Supplementary Figure S2), and iv) it has been previously reported to play a role in the maintenance of an open chromatin state^28–30^. For all these reasons and to further dissect the role of miR-144 in chromatin regulation, we decided to focus from here forward on Hmgn2.

### Hmgn2 is a conserved target of miR-144

*Pre-miR-144* can generate two mature isoforms that are variable at their 5’ ends by 1 nucleotide due to the reshaping of its terminal loop^31^. According to TargetScanFish v6.2, *Hmgn2* mRNA from zebrafish (*Danio rerio*) has two predicted 7mer-m8 target sites in for miR-144-v1 its 3’UTR^32^ (Figure 2E and Supplementary Figure S2). Moreover, the expression of Hmgn2 is stabilized in miR-144^Δ/Δ^ mutants (Figure 2A). To experimentally demonstrate that *Hmgn2* is a direct target of miR-144, we injected a Nanoluciferase reporter fused to the zebrafish *Hmgn2* 3’UTR (Nluc-*Hmgn2*-3’UTR-wt) into zebrafish embryos. Only the co-injection of the reporter with miR-144 duplex repressed reporter expression (Figure 2F). Conversely, mutation of the miR-144 target site in the 3’UTR of *Hmgn2* disrupted miR-144-mediated repression of the Nanoluciferase reporter (Figure 2F).

To probe the expression domain of endogenous Hmgn2 and its regulation by miR-144, we detected the expression of *Hmgn2* mRNA in 30-hpf embryos using *in situ* hybridization. It revealed a distinct pattern of Hmgn2 expression in the notochord. In addition, expression of Hmgn2 was detected in the posterior blood island, an early hematopoietic site of the embryo, in miR-144^Δ/Δ^ but not in wild-type siblings^33^ (Figure 2D). This observation can be explained by the stabilization of the *Hmgn2* transcript in the miR-144 mutant embryos. This result demonstrates that miR-144-mediates repression of the endogenous *Hmgn2* in the hematopoietic tissue and suggests that *Hmgn2* down-regulation occurs early in the erythrocyte maturation process.

To gain information about Hmgn2 regulation dynamics during erythropoiesis, we conducted single-cell RNA-Seq (scRNA-Seq) analysis of the wild-type and miR-144^Δ/Δ^ pronephros, the main hematopoietic organ of adult fish^34^ (Supplementary Figure S3). Focusing on the erythroid lineage, we observed that miR-144^ι1/ι1^ erythroid cells do not reach their final differentiation state and accumulate at intermediate stages when compared to wild-type cells (Figure 2G). The transcriptome composition of miR-144^ι1/ι1^ cells resembles intermediate differentiation stages of wild-type erythroid progenitors. This fact indicates that the loss of miR-144 significantly disrupts the terminal differentiation process blocking the maturation of erythroblasts.

When we follow the expression of Hmgn2 in the developmental trajectory from the hematopoietic progenitor cell cluster to the cluster of mature erythrocytes, we observed sustained expression of Hmgn2 in miR-144^ι1/ι1^ cells while the expression of Hmgn2 was reversely correlated with differentiation progression in wild-type siblings (Figure 2G). It demonstrates that loss of miR-144 leads to increased expression of Hmgn2 in its endogenous context and correlates with impaired differentiation progression. These results indicate that miR-144 is involved in the regulation of Hmgn2 expression *in vivo* at least in part through seed-mediated interactions in its 3’UTR that are conserved across vertebrates.

### Disruption of miR-144-mediated regulation of Hmgn2 mimics the erythropoietic defects of the miR-144 mutant

To evaluate the importance of the miR-144/Hmgn2 regulatory axis *in vivo*, we genetically probed this regulatory circuit. We aimed to disrupt the direct interaction between miR-144 and Hmgn2 while preserving their expression in their cell-specific context. First, we attempted to delete the two miR-144 target sites in the endogenous Hmgn2 3’UTR of zebrafish. However, the lack of appropriately positioned guide RNAs in this genomic region precluded us from using a CRISPR/Cas9 system and instead drove us to apply a transgenic approach. To recapitulate the endogenous regulation of Hmgn2 by miR-144 we devised a transgenic zebrafish line expressing coding regions of Hmgn2 fused to the full-length *Hmgn2* 3’UTR that contains miR-144 target sites (Figure 2E). An EYFP sequence was included in this construct to monitor the expression of the transgene and was separated from Hmgn2 by a T2A site. The 3’UTR comprising miR-144 target sites was flanked by *loxP* sites that could be used to delete by recombination the miR-144 sites of the transgene cassette and generate a control transgenic line in which miR-144-mediated regulation of Hmgn2 is disrupted (Figure 3C and 3D).

**Figure 3.**
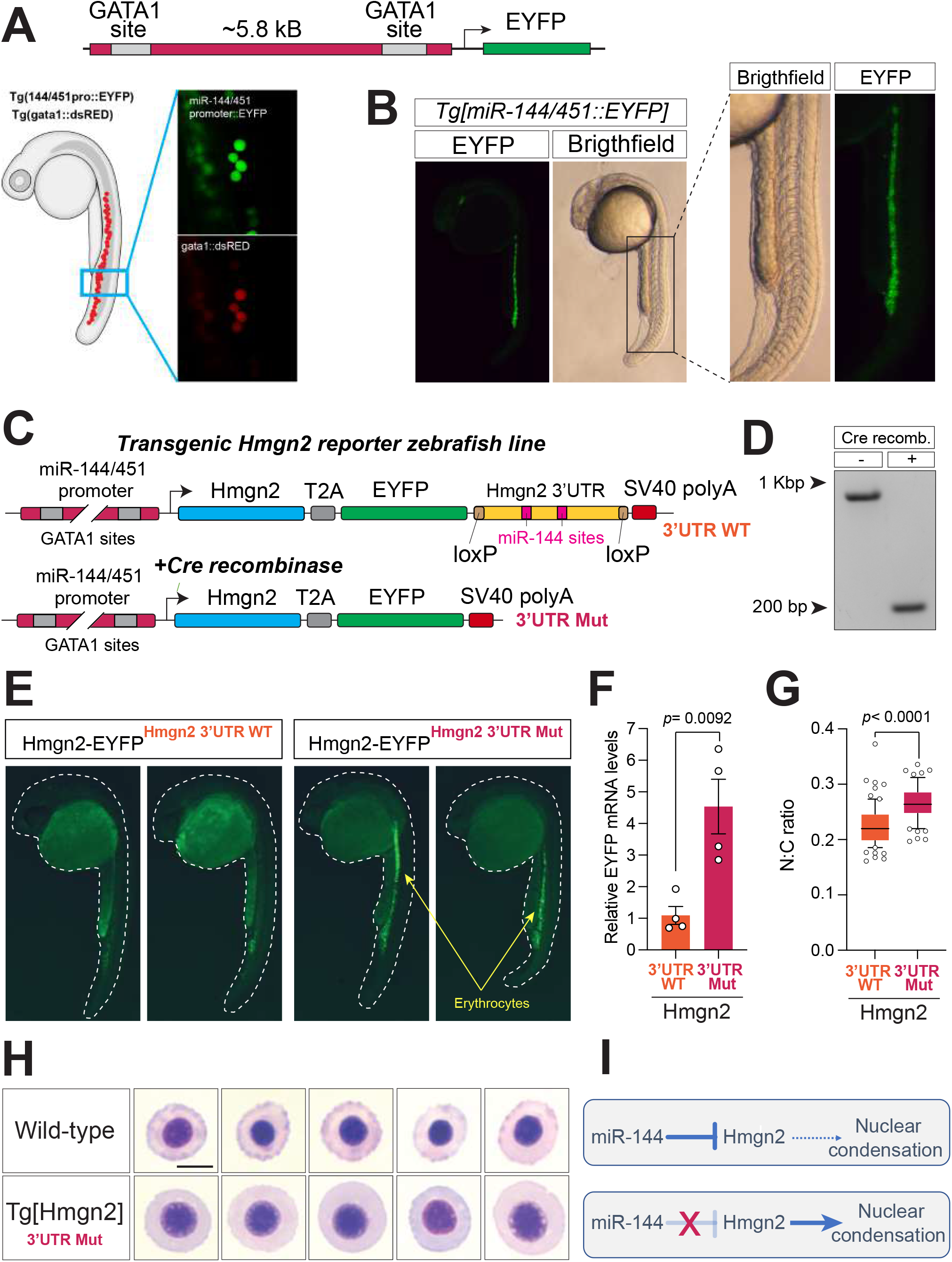
**(A)** A schematic representation of a transgene expressing EYFP from miR-144/451 promoter and its validation by colocalization with dsRed that is expressed from gata1 promoter in double transgenic fish. Photos are taken at 24 hpf. **(B)** miR-144/451 promoter drives the expression of EYFP in the posterior blood island and caudal vein. **(C)** A schematic representation describing a strategy of generating a transgenic line to analyze the expression of Hmgn2 in zebrafish. **(D)** PCR validation of efficient 3’UTR deletion by Cre-mediated recombination. **(E)** EYFP expression in 24 hpf old embryos with (right) and without (left) *Cre* mRNA injection. **(F)** Real-time quantitative PCR of Hmgn2-EYFP transgene in control and injected with *Cre* mRNA embryos at 24 hpf. Expression is normalized *GAPDH* mRNA. Data represent mean ± standard error of the mean of four technical replicates. *p*-values from unpaired *t-*test. **(G)** Quantitative analysis of the N:C ratio of erythrocytes stained in (H). Between 66 and 70 cells are analyzed in each case. *p*-values from unpaired *t-*test. Boxes enclose 10 to 90 percentiles. **(H)** May-Grunwald-Giemsa staining of peripheral blood cells isolated from the transgenic embryos with mutant Hmgn2 3’UTR and wild-type siblings at 2-dpf. **(I)** A schematic describing miR-144/Hmgn2 regulatory axis.

To ensure that the Hmgn2-EYFP transgene has the same spatial and temporal expression as miR-144, we cloned and used the miR-144/451 promoter to drive the expression of the transgene (Figure 3A). We defined the miR-144/451 promoter as the 5.4 Kb region upstream of the start of transcription of the miR-144/451 cluster that encompasses two GATA1 binding sites that were characterized previously^6^. First, we confirmed that this promoter drives the expression of a fluorescence reporter gene (EYFP) specifically in the erythrocytes. When we crossed the *miR-144/451::EYFP* line to the *gata1::dsRed* line we detected a perfect overlap in the expression of EYFP and dsRed (Figure 3A). We used this validated promoter to drive the expression of the aforementioned transgenic construct *miR-144/451::Hmgn2-EYFP-3’UTR-WT*. We observed that the expression of EYFP was barely detectable at 24-hpf in the wild-type background (Figure 3E). This could be explained by repressive elements present in its 3’UTR, such as miR-144 target sites. To test this hypothesis, we injected mRNA encoding Cre recombinase in part of the F1 generation off-spring of *miR-144/451::Hmgn2-EYFP-3’UTR-WT* to generate the *miR-144/451::Hmgn2-EYFP-3’UTR-MUT* line that carries a truncated 3’UTR without the miR-144 target sites. Cre-mediated recombination of the transgene is highly efficient, as shown by PCR of the transgene 3’UTR, leading to 100% efficiency of Hmgn2 3’UTR deletion (Figure 3D). This experimental design allows us to compare two zebrafish lines with different 3’UTRs avoiding variability due to positional effects caused by independent integration of the transgene cassette at different genomic regions. Live imaging of transgenic embryos at 1-dpf revealed that *miR-144/451::Hmgn2-EYFP-3’UTR-MUT* embryos express significantly higher levels of EYFP in the posterior blood island and caudal vein when compared to transgenic embryos carrying the wild-type *Hmgn2* 3’UTR (Figure 3E). These results indicate that Hmgn2 expression is regulated *in vivo* in the erythrocytes by *cis* elements contained in its 3’UTR, which include the miR-144 sites.

To gain quantitative information about Hmgn2 transgene expression we performed RT-qPCR analysis. We compared the expression of Hmgn2-T2A-EYFP with and without the *Hmgn2* 3’UTR. We observed that Cre-mediated deletion of the 3’UTR leads to over 4-fold up-regulation of transgene expression (Figure 3F). This observation indicates that miR-144 is a strong repressor of Hmgn2 expression in erythrocytes and validates the observation obtained by fluorescent microscopy (Figure 3E). Nevertheless, we cannot exclude that additional factors may also contribute to *Hmgn2* mRNA down-regulation.

We decided to evaluate the consequences of Hmgn2 upregulation on erythrocyte morphology. We isolated peripheral blood cells from the *miR-144/451::Hmgn2-EYFP-3’UTR-MUT* in a wild-type background, which expresses miR-144. We observed that erythrocytes over-expressing Hmgn2 displayed a higher N:C ratio as compared to erythrocytes from wild-type zebrafish (Figure 3G and H). Altogether, these results provide a genetical validation that post-transcriptional regulation of Hmgn2 is necessary for proper erythrocyte maturation and that the disruption of this regulation can mimic the loss-of-miR-144 phenotype (Figure 3I).

### Loss of function of Hmgn2 rescues the erythropoietic defects of miR-144 mutant

To counterbalance the previous experiments, we set out to rescue the miR-144 mutant genetically. We hypothesized that reducing the dosage of Hmgn2 in the miR-144^ΔΔ^ background would compensate for the lack of post-transcriptional down-regulation of Hmgn2 to an extent that may restore normal erythrocyte maturation. To this end, we generated a 75-nucleotide deletion spanning intron 1 and exon 2 of Hmgn2 using CRISPR/Cas9 (Figure 4A). According to high-throughput sequencing data, this deletion still allows for the in-frame expression of Hmgn2 skipping exon 2, which that deletes 12 out of 75 amino acids (Supplementary Figure S4A). This Hmgn2 variant appears to be non-functional because erythrocytes from homozygous maternal-and-zygotic *Hmgn2* mutant (Hmgn2^ΔΔ^) embryos show premature nuclear condensation compared to wild-type siblings (Figure 4C). These results suggest that our Hmgn2^Δ/Δ^ is at least a partial loss-of-function mutant and indicate that Hmgn2 is necessary to maintain proper nuclear organization during erythropoiesis. When we performed ATAC-Seq on erythrocytes isolated from Hmgn2^Δ/Δ^ we observed a decrease in open chromatin regions which opposes the phenotype observed in the miR-144 mutant (Figure 4D and 1H). Altogether these results confirm the role of Hmgn2 as a factor necessary to maintain an open chromatin state. Next, we crossed Hmgn2^Δ/Δ^ mutant fish with miR-144^Δ/Δ^ to generate the double-mutant zebrafish line miR-144^Δ/Δ^/Hmgn2^Δ/Δ^. When we probed erythrocyte development in 2-dpf embryos, we observed that the N:C ratio of erythrocytes of this double mutant is similar to the ratio from wild-type siblings (Figure 4F), indicating that reduced Hmgn2 activity rescues the effect of the loss miR-144 and suggesting that Hmgn2 is an important player in driving the condensation defects of the miR-144 mutant. However, if Hmgn2 were the only factor driving the N:C ratio, we would expect that the N:C ratio of the double mutant would mimic that of single the Hmgn2 mutant due to the inability to maintain open chromatin state even before miR-144 is expressed. Since this is not the case, these results suggest that other factors that are stabilized in the miR-144 mutant regulate the N:C ratio of the cells (Figure 2A). Unexpectedly, the rescue of nuclear condensation defects of the double mutant (Figure 4F) does not translate into changes in chromatin accessibility analyzed by ATAC-seq (Supplementary Figure S4B), indicating that the morphological rescue (Figure 4F) is driven by additional mechanisms triggered by the loss of miR-144.

**Figure 4.**
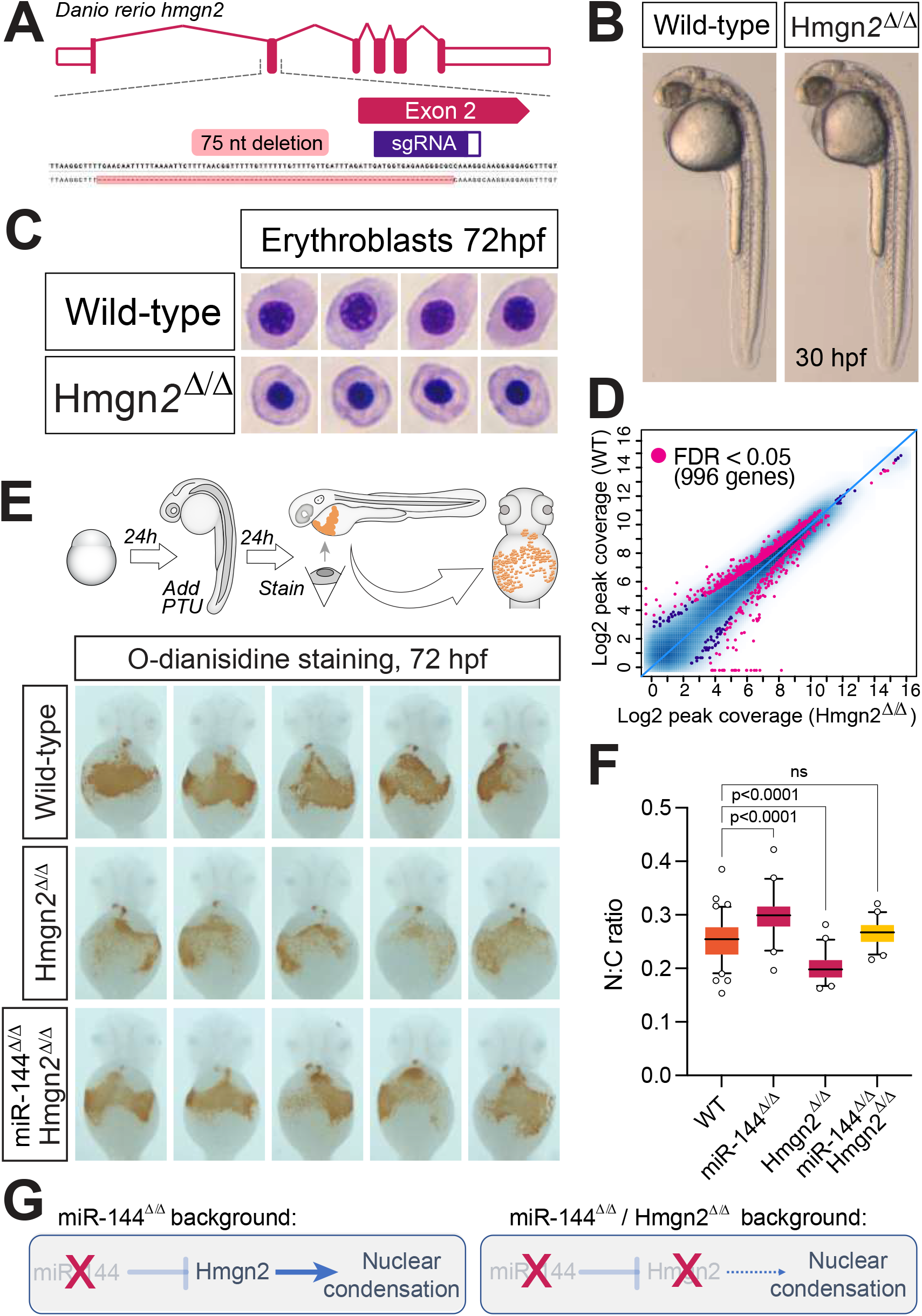
**(A)** Schematic representation of Hmgn2 gene from Danio rerio and CRISPR/Cas9 induced deletion. **(B)** Maternal-zygotic Hmgn2 (Hmgn2^Δ/Δ^) mutants at 30 hpf. **(C)** May-Grunwald-Giemsa staining of peripheral blood cells isolated from Hmgn2^Δ/Δ^ and wild-type siblings at 2 dpf. **(D)** Scatter plot showing differential chromatin accessibility in Hmgn2^Δ/Δ^ and wild-type using ATAC-Seq. Each dot represents an ATAC-seq peak. **(E)** *O*-dianisidine staining of 2-dpf embryos treated with PTU to reveal hemoglobinized cells. **(F)** Quantitative analysis of nucleocytoplasmic ratio of erythrocytes isolated from miR-144^Δ/Δ^, miR-144^Δ/Δ^/Hmgn2^Δ/Δ^ and wild-type sibling stained with May-Grunwald-Giemsa to rescue miR-144^Δ/Δ^ phenotype. Between 42 and 93 cells are analyzed in each case. p-values from ordinary one-way ANOVA. Boxes enclose 5 to 95 percentiles. **(G)** Schematic of genetic probing of miR-144/Hmgn2 regulatory axis.

*O*-dianisidine staining revealed that Hmgn2^Δ/Δ^ embryos display mild anemia under oxidative stress conditions compared to wild-type embryos (Figure 4E). This defect is compensated for by the additional loss of miR-144 (Figure 4E), probably through the stabilization of other chromatin factors described before and is in agreement with the rescue of the erythrocyte N:C ratio (Figure 4F). Altogether, these results suggest that reducing Hmgn2 levels prevents the formation of the phenotype induced by the loss of miR-144 and re-establishes normal erythrocyte maturation (Figure 4G).

### miR-144/Hmgn2 regulatory axis facilitates differentiation of human erythroid progenitor cells

To test the relevance of the miR-144/Hmgn2 axis during human erythropoiesis, we took advantage of a cell derived from human induced pluripotent stem cells (iPSCs) that can recapitulate erythroid differentiation *in vitro*. These iPCS-derived erythroid progenitors represent cells captured at a proerythroblast stage that can undergo further erythroid differentiation according to the previously established protocol^35^. First, we analyzed expression of miR-144 during the time course of differentiation and observed a significant up-regulation of its expression, as well as of its cluster neighbor, miR-451 (Figure 5C).

**Figure 5.**
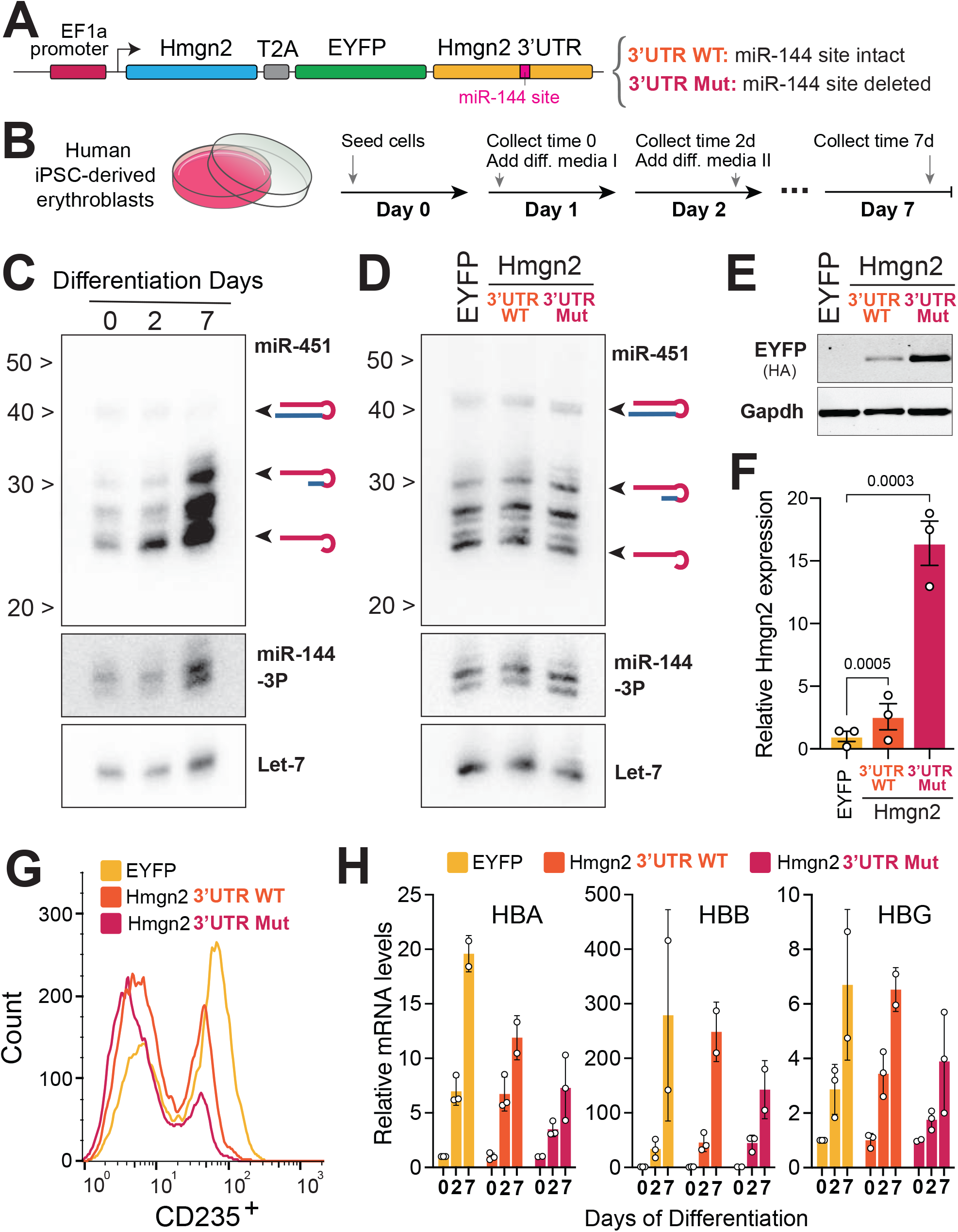
**(A)** A schematic representation describing a strategy of generating cell lines to analyze the expression of Hmgn2 in human iPSC-derived erythroblast cells. **(B)** Experimental protocol to further induce the differentiation of human iPSC-derived erythroblast cells. **(C)** Northern blot analysis to detect miR-451, miR-144 and let-7 during the time course of cell differentiation. **(D)** Northern blot analysis to validate that over-expression of transgenic construct does not affect miR-451 and miR-144 levels at the time point 0. **(E)** Western blot to detect accumulation of EYFP which is expressed from the transgenic construct either with WT UTR or with mutant miR-144 target site at the time point 0. **(F)** Real-time quantitative PCR of Hmgn2-EYFP transgene in at the time point 0. Expression is normalized on *GAPDH* mRNA. Data represent mean ± standard error of the mean of three technical replicates. *p*-values from ordinary one-way ANOVA. **(G)** Flow cytometry analysis (FACS) of erythroid cells at time point 2 to quantify the levels of CD235. **(H)** Real-time quantitative PCR of globin A (HBA) and globin E (HBE) at indicate collection time points. Expression is normalized on *GAPDH* mRNA.

To test the effect of miR-144 on Hmgn2 expression in human cells and its importance for erythroid differentiation, we applied a similar strategy as we used in zebrafish. We uncoupled Hmgn2 from miR-144-mediated regulation by expressing Hmgn2 from a lentivirus construct encoding either the wild-type human Hmgn2 3’UTR (*hs*Hmgn2-EYFP-3’UTR-WT) or the 3’UTR with a small deletion removing miR-144 target site (*hs*Hmgn2-EYFP-3’UTR-MUT) (Figure 5A). As a control, we infected erythroid cells with a lentivirus expressing EYFP alone. Cells expressing the construct with wild-type Hmgn2 3’UTR display a modest increase in Hmgn2-EYFP expression (2-fold over control), while cells expressing Hmgn2 with mutant Hmgn2 3’UTR expressed Hmgn2 15-fold higher over the control (Figure 5E and 5F). We confirmed that the expression of EGFP or Hmgn2 did not interfere with the levels of miR-144 in cells expressing lentiviral constructs, suggesting that the difference in the expression of Hmgn2-EYFP is not due to different levels of miR-144 in these cells (Figure 5D), but would rather be caused by the deleted target site in its 3’UTR. Next, all three engineered lines were exposed to maturation media to induce further differentiation. We evaluated expression of several markers at different time points to determine the differentiation progress of each cell line. We measured the expression of CD235 (Glycophorin A), a hallmark of erythroid specific lineage^36^. We detected the expression of CD235 by flow cytometry in all samples at the end of the differentiation regime (day 7). However, the CD235 levels were inversely correlated with Hmgn2 expression (Figure 5G). We also quantified the expression of globins at 2 and 7 days after the start of differentiation (Figure 5H). We observed that the onset of globin HBA, HBB and HBG expression was delayed and reduced in cells expressing the *hs*Hmgn2-EYFP-3’UTR-MUT (Figure 5H). This phenotype was milder in cells expressing the wild-type 3’UTR. We observed that cells expressing more Hmgn2 do not reach the same level of globin expression as cells with lower levels of Hmgn2, which may indicate their reduced competence for differentiation. Altogether, these results suggest that the miR-144/Hmgn2 regulatory axis is functional in humans and required for proper erythropoiesis.

## DISCUSSION

Here we report the unique finding that miR-144 single-handedly regulates multiple chromatin factors to force erythrocyte progenitors into the differentiation pathway. In particular, we identified and genetically probed how a new miR-144/Hmgn2 regulatory axis governs nuclear condensation during erythropoiesis. Hmgn2 is a chromatin regulator that binds to nucleosomes and maintains chromatin in a transcriptionally active state^28,29,37,38^. Our data show that sustained expression of Hmgn2 in the erythrocyte lineage after deletion of miR-144 target sites in its 3’UTR mimics the miR-144 mutant phenotype in zebrafish erythroblasts and human iPSC-derived erythroid cells. Conversely, we demonstrate that disruption of Hmgn2 activity partially rescues the nuclear defects caused by the loss of miR-144. These results clearly intersect with previous findings showing that Hmgn2 preferentially binds to chromatin regulatory sites, including the binding to mononucleosomes containing the adult globin gene cluster^39^ and its role as a transcriptional activator^40^. Our data expands the impact of miR-144 activity well beyond its seed-matching target mRNAs and illustrate the broad impact of miRNA activity in shaping global gene expression. Ultimately, the significance of these results is that they challenge the current dogma in the field that perceives chromatin-modulating microRNAs as gatekeepers of cell identity and inhibitors of erythrocyte differentiation^7,41,42^. Instead, we present compelling evidence that miR-144 plays a pivotal role in enhancing the process of erythrocyte maturation from fish to humans. This paradigm shift not only enriches our understanding of miRNA-mediated regulatory mechanisms during development but also will open new therapeutic opportunities to treat erythropoiesis-related disorders.

While the downregulation of multiple miRNAs has been implicated in the post-transcriptional regulation of erythrocyte maturation, here we present an opposite example whereby the activity of the miRNA is necessary to proceed through the erythrocyte maturation path. For instance, decreasing miR-191 levels are necessary to up-regulate the expression of Riok3 and Mxi1, which in turn antagonize the activity of histone acetyltransferase Gcn5 and facilitate chromatin condensation^7^. Similarly, miR-181, miR-30a, miR-34a, and miR-9 suppress chromatin condensation, and their down-regulation is required for normal erythrocyte maturation and nuclear extrusion^41,42^. Contrary to these known examples, the expression of miR-144 during erythropoiesis is necessary to induce nuclear condensation and subdue transcription. This previously unappreciated role of miR-144 in chromatin regulation neatly complements the function of miR-144 that we recently described as a master repressor of canonical microRNAs via its direct targeting of *Dicer* mRNA in erythrocytes^17^. Altogether, it presents a picture whereby the combined action of multiple miRNAs regulates a cell’s entry into the erythrocyte differentiation pathway. It is the role of miR-144 to simultaneously downregulate these gatekeeper miRNAs by targeting Dicer^17^ and to collaborate with other parallel mechanisms^43^ to repress Hmgn2, thereby inducing erythrocyte maturation via chromatin remodeling.

Erythrocyte maturation is heavily regulated at the chromatin level as nuclear condensation is an essential part of the process. In mammals, this process culminates with the extrusion of the nucleus^44^. A global decrease in histone acetylation (such as H3K9Ac, H4K5Ac, H4K12Ac, and H4K8Ac) and increased methylation (H3K79 and H4K20) have been reported to be important for normal chromatin condensation^45–47^. In addition to epigenetic changes, the release of histones mediated by caspase-3 cleavage and ubiquitin-dependent degradation of nuclear lamin B^48,49^ is also required for proper nuclear compaction during erythropoiesis. In the current work, we identify another regulatory mechanism that contributes to chromatin condensation during erythropoiesis. We demonstrate that repression of Hmgn2 and other chromatin factors allows chromatin condensation to proceed. High-mobility group nuclear proteins are a family of highly abundant non-histone chromatin binding proteins that modulate chromatin organization^30,50^. Among them, proteins of the HMGN family specifically remodel chromatin by antagonizing the binding of linker histone H1 to nucleosomes near chromatin regulatory sites^29,30^. From this vantage point, HMGN proteins play an important role in development. Their expression is high in mouse embryonic stem cells (ESCs)^51^ and iPSCs^29^, but gradually decays during tissue differentiation^52^. This phenomenon has been described for the development of multiple tissues, including the eye^53,54^, hair follicle^55^ and during myogenesis^56^, chondrocyte differentiation^57^, oligodendrocyte lineage specification^51^, and erythropoiesis^58^. The expression dynamics of HMGN proteins suggest that their collective activity is necessary to maintain stem cell identity^28^ but it must be dampened to proceed with cell differentiation. While changes in the chromatin structure of the HMGN genes during erythrocyte differentiation reduce in part the expression of these genes^43^, here we present a parallel mechanism that down-regulates Hmgn2 expression post-transcriptionally. Future work will establish if the regulation of Hmgn2 expands to other members of the high mobility group nuclear protein family regulated by miR-144 or other miRNAs as a prevalent regulatory mode during the differentiation of other tissues.

## ACKNOWLEDGMENTS

We thank A. Grishok, X. Varelas, M. Blower, M. Garcia-Marcos and L. Zon for fruitful discussions and access to instruments and microscopes. We thank Yuriy Alekseyev, Joshua Campbell and the Boston University Microarray and Sequencing Resource for conducting sRNA-Seq and scRNA-Seq sequencing and analysis. We thank Nicki Watson (Harvard) for conducting Transmission Electron Microscopy. We thank Boston University Flow Cytometry Core for cell sorting. We thank Charith Wijeyesekera for the help with iPSCs experiments. We thank Andrew Tilston-Lunel for taking photos of *miR-144/451::EYFP/gata1::dsRed* transgenic. This work was supported by the National Institute of Health (US) R01GM130935-03, Boston University Genome Science Institute scRNA-Seq seed grant and RNA-sequencing award.

## AUTHOR CONTRIBUTIONS

D.A.K. designed, performed and analyzed the experiments and wrote the draft of the manuscript. L.F. analyzed ATAC-sequencing data. A.M.M. performed whole mount *in situ* hybridization. N.S. cloned candidate genes and helped with generation and maintenance of zebrafish lines. I.A.W. performed sample preparation for single cell analysis of zebrafish pronephros. S.M. analyzed Quant-sequencing, RNA-sequencing data and single-cell sequencing data. K.V. and G.M. designed and helped with human iPSCs experiments and erythroid differentiation. D.C. conceived and supervised the project, performed experiments, analyzed the data, acquired funding and wrote the manuscript.

## METHODS

### Zebrafish strains

Zebrafish strains were bred, handled, and maintained according to the standard laboratory conditions under IACUC protocol PROTO201800373 at Boston University. Experiments were performed in hybrid wild-type strain crosses obtained from AB/TU and TL/NIHGRI breeders.

### Generation of mutant zebrafish lines using CRISPR/Cas9

miR-144 mutant line (miR-144^Δ/Δ^) was generated and described previously in our laboratory^17^. To generate Hmgn2^Δ/Δ^ we designed sgRNA targeting exon 2 of *hmgn2* gene was designed using CRISPRscan^59^ yielding the following target sequence: 5’-GATGGTGAGAAGGGCGCCAA(AGG)-3’; where the Cas9 PAM sequence (NGG) is between parentheses. sgRNA templates were generated by annealing and polymerase-mediated extension of a forward oligo containing the T7 promoter sequence, the 20 nt sgRNA target sequence (without the PAM sequence) and a 15 nt sequence complementary to the reverse oligo containing the invariable Cas9-binding scaffold. PCR reactions with Q5 high fidelity polymerase (NEB) were carried out as follows: 1 cycle 95°C for 3 minutes; 35 cycles (95°C for 3 minutes, 55°C for 30 s, 72°C for 20 s); 1 cycle at 72°C for 5 minutes. Reactions were purified with a PCR purification kit (NEB). Approximately 120–150 ng of DNA was used as a template for a T7 *in vitro* transcription (IVT) reaction with the AmpliScribe-T7-Flash transcription kit (Epicenter ASF3507). IVT sgRNA products were purified Monarch PCR and DNA Cleanup Kit (NEB) and quantified. Zebrafish embryos were injected at one-cell stage with *Cas9* mRNA (100 pg) together with 30 pg of sgRNA to generate *hmgn2* mutant. miR-144^Δ/Δ^/Hmgn2^Δ/Δ^ mutant was generated by crossing miR-144^Δ/Δ^ to Hmgn2^Δ/Δ^ line, followed by genotyping and subsequent in-crossing until obtaining the miR-144^Δ/Δ^/Hmgn2^Δ/Δ^ fish line.

### Generation of transgenic zebrafish lines

To generate transgenic zebrafish line expressing a fluorescent EYFP reporter in erythrocytes we first cloned the miR-144/451 promoter, the 5.4 kb region located upstream of miR-144/451 locus. We cloned pDEST-miR-144/451-EYFP plasmid from p5E-miR-144/451 promoter, pME-EYFP and p3E-polyA plasmids using Gateway LR Clonase II Enzyme Mix (Invitrogen). We generated *miR-144/451::EYFP* transgenic line using Tol2 transposon system (REF). To validate miR-144/451 promoter *miR-144/451::EYFP* line was crossed *gata1::dsRed* line (Ref).

Then we generated pDEST-miR-144/451-*dre*Hmgn2-EYFP-*dre*Hmgn2-3’UTR-WT plasmid from p5E-miR-144/451 promoter, pME-*dre*Hmgn2-EYFP and p3E-*dre*Hmgn2-3’UTR-WT-polyA using Gateway LR Clonase II Enzyme Mix (Invitrogen). We included two loxP site surrounding to *dre*Hmgn2-3’UTR-WT. We generated miR-144/451-*dre*Hmgn2-EYFP-*dre*Hmgn2-3’UTR-WT line using Tol2 transposon system. In order to generate *miR-144/451::Hmgn2-EYFP-3’UTR-MUT* line, lacking most of it 3’UTR including miR-144 target sites, we injected mRNA encoding Cre recombinase (1 nL of 5 ng/mL) and validated 3’UTR removal by PCR.

### Transmission Electron Microscopy

The cells were fixed in 2.5% gluteraldehyde, 3% paraformaldehyde with 5% sucrose in 0.1M sodium cacodylate buffer (pH 7.4), pelleted, and post fixed in 1% OsO_4_ in veronal-acetate buffer. The cells were stained en block overnight with 0.5% uranyl acetate in veronal-acetate buffer (pH 6.0), then dehydrated and embedded in Embed-812 resin. Sections were cut on a Leica ultra-microtome with a Diatome diamond knife at a thickness setting of 50 nm, stained with 2% uranyl acetate, and lead citrate. The sections were examined using a Hitachi 7800 and photographed with an AMT ccd camera. To quantify euchromatin content of nuclei we used ImageJ software. First, images were converted to 8-bit format, then nuclear area was manually selected and threshold was adjusted (using IsoData auto setting).

### Immunofluorescent Microscopy

Erythrocytes were isolated from 3-dpf zebrafish embryos (wild-type and miR-144^Δ/Δ^) by dissecting their tail using sapphire blade (WPI) and collecting cells in the bleeding buffer (1X PBS containing 2% PBS and 5 mM EDTA). Erythrocytes were washed twice with the bleeding buffer and spread on Superfrost Plus microscope slides using Shandon Cytospin 2 Centrifuge and fixed for 20 min in ice-cold methanol. Erythrocytes were incubated for 30 min at room temperature in blocking buffer and stained directly on slides with anti-RNAP II Phosho-Ser2 (Abcam #5095) in the blocking buffer (1X PBS, bovine serum albumin (BSA) 2%, Tween 0.2%) overnight at 4°C. The slides were then washed three times with 1X PBS, after what anti-rabbit Alexa Fluor 488 secondary antibodies (Jackson Immuno Research Laboratories, #711-545-152) in blocking buffer were added and cells were incubated for 1h at room temperature. After that DAPI was added to cells and incubated for 10 min at room temperature. After final washes with 1X PBS and mounted using Antifade Mounting Media (Vestashield) for fluorescence microscopy imaging. Images were captured using Zeiss Axio Observer Z1 microscope equipped with a digital camera (C10600/ORCA-R2 Hamamatsu Photonics).

### May-Grunwald Giemsa staining

Peripheral blood from different developmental timepoints was isolated from wild-type, miR-144^Δ/Δ^, Hmgn2^Δ/Δ^, miR-144^Δ/Δ^/ Hmgn2^Δ/Δ^ and Tg::mR-144/451:Hmgn2 zebrafish by dissecting their tail using sapphire blade (WPI) and collecting cells in the bleeding buffer (1X PBS containing 2% PBS and 5 mM EDTA). Erythrocytes were spread on Superfrost Plus microscope slides using Shandon Cytospin2 at 500 rpm for 5 min. Then cells were fixed in iced cold methanol for 5 min and air-dried. Cells were incubated for 5 min in May-Grunwald Stain (Sigma). After that slides were rinsed twice in 1X PBS in transferred into dilute Giemsa solution (1:20) (Sigma) for 20 min. Finally, briefly rinsed in deionized water and air-dried (Sigma). Images were captured using a Zeiss Axio Observer.Z1 microscope with Zeiss ZEN 3.3 Blue software.

### *O*-dianisidine staining

Hemoglobin was detected in 2-dpf embryos by incubation for 15 min in the dark in *O*-dianisidine staining solution (2 mL of water, 2 mL of 0.7 mg/mL *O*-dianisidine, dissolved in 96% ethanol and protected from light, 0.5 mL of 100 mM sodium acetate, 100 mL of 30% hydrogen peroxide) and then transferred into 1X PBS. To apply oxidative stress, live embryos were transferred to water containing 0.003% phenylthiourea (PTU) from 8 hpf until collection time at 2-dpf.

### Whole-mount *in situ* hybridization

Expression of endogenous hmgn2 was detected by whole mount *in situ* hybridization, following the protocols described in Thisse C. *et al*^60^. Briefly, a 792-bp segment of hmgn2 cDNA was amplified by PCR with oligos that also added a T7 promoter to the amplicon. After agarose gel purification of the PCR amplicon, it was in vitro transcribed with the T7 transcription kit (Invitrogen #AM1333) and the DIG labeling RNA mix/kit (Roche #39354521) to generate a DIG-labeled antisense probe, complementary to *hmgn2* mRNA. 1-day-old embryos where fixed with 4% paraformaldehyde (PFA) solution in PBS at pH 7 overnight. After fixation, the embryos were gradually dehydrated in 100% methanol and stored at −20°C until being used. To initiate the *in situ* hybridization, embryos are gradually rehydrated with PBS pH 7, and permeabilized by proteinase K digestion (10 µg/ml) at room temperature for 10 minutes. Proteinase K digestion was stopped by fixation with 4% PFA. Pre-hybridization and onvernigth hybridization with the DIG-labled RNA probe was performed as described previously^60^. After washing and blocking, embryos were incubated with an anti-DIG antibody conjugated to alkaline phosphatase in blocking buffer overnigth at 4°C. Finally, embryos are developed with the staining solution containing NBT and BCIP. Embryos were transferred to a stop sulition once the apporpiate color intensity was detected. Embryos were imaged in a stereomicroscope Stemi 508, using the Zen suite (Zeiss). Full details of washing steps and buffer compositions can be found in ^60^.

### ATAC-sequencing

We followed the protocol from Yang, S. et al ^61^ with some modifications. Erythrocytes were isolated from WT and miR-144^Δ/Δ^ adult zebrafish. Peripheral blood was collected from cut fin of adult fish, or from 2-dpf zebrafish embryos by dissection of caudal vein using sapphire blade, in the collection buffer (1X PBS containing 2% PBS and 5 mM EDTA). Number of blood cells was quantified with Countess Cell Counting Chamber Slides (ThermoFisher) and 50,000 cells were used for every reaction. Collected cells were spun down in the cell collection medium at 500 x g for 5 min at 4°C. Cell pellets were gently washed with 50 μL of cold 1X PBS and spun down at 500 g for 5 min at 4°C. After removal of PBS, 50 μL of cold cell lysis buffer (10 mM Tris–HCl pH 7.4, 10 mM NaCl, 3 mM MgCl2, 0.1% IGEPAL CA-360) was added, and cells were resuspended by gentle pipetting followed by an immediate spin at 500 g for 5 min at 4°C. The supernatant was removed and the purified nuclei were resuspended in the transposition reaction mixture (25 μl 2X TD Buffer, 2.5 μl Tn5 transposase, 22.5 μl Nuclease Free water) and incubated for 90 minutes at 37°C. DNA was then purified with Monarch PCR & DNA Cleanup Kit (New England BioLabs). Libraries were prepared using Q5 High-Fidelity 2X Master Mix NEB, M0492) with the following conditions: 72°C, 5 minutes; 98°C, 30 seconds; 15 cycles of 98°C, 10 seconds; 63°C, 30 seconds; and 72°C, 1 minute. Amplified libraries were purified with Monarch PCR & DNA Cleanup Kit (New England BioLabs) and sequences Illumina NextSeq 2000 system at Boston University Microarray and Sequencing core. ATAC-Seq data was analyzed as described in Yang, S. et al ^61^.

### Analysis of candidate genes on erythrocyte morphology

To analyze the effect of over-expression candidate genes (*hmgn2, gtf2a1, cbx8a, nap1l4b and dicer1*) their mRNAs were injected into 1-cell stage wild-type zebrafish embryos (100 pg per embryo). 2-dpf peripheral blood was isolated as described above and analyzed by May-Grunwald Giemsa staining.

### MicroRNA reporter assay

Full length *hmgn2* 3’ UTR from zebrafish and its variant lacking two miR-144 targets sites were cloned into pCS2+ after Nanoluciferase coding sequence. Reporter constructs were linearized with NotI restriction enzyme and *in vitro* transcribed with mMESSAGE mMACHINE SP6 Transcription Kit (Ambion). For the fluorescent miRNA reporter assay, zebrafish embryos were injected with 1 nL of 100 ng/mL of WT 3’UTR *hmgn2* or MUT 3’UTR *hmgn2* reporter together with firefly luciferase^62^ as a control reporter. Synthetic RNA oligonucleotides (IDT) representing miR-144 duplex were annealed by incubation in 1X TE buffer at 90 °C for 5 min and then slow cooled to room temperature. 1 nL of 10 mM miR-144 duplex was injected together with reporters. To quantify the activity of reports groups of 5 embryos were collected 8 hours post-injection in triplicates and lysed in 100 mL of lysis buffer (Promega). Reporter expression was quantified with the Nano-Glo Dual-Luciferase Reporter Assay System (Promega) in a Synergy H1 Hybrid Multi-Mode Microplate Reader (Thermo).

### Gene expression analysis using RNA-Seq

Peripheral blood cells were isolated from WT and miR-144^Δ/Δ^ zebrafish at 3-dpf in biological duplicates. Total RNA was extracted with TRIzol and quantified. Poly(A) mRNA selection and library preparation was performed using NEBNext Ultra II RNA Kit (NEB). Sequencing was performed at Boston University Microarray and Sequencing core. Sequencing data was analyzed as follows: Transcript based counts were obtained from Kallisto (v0.46.1) using *Danio rerio* GRCz10 cDNA annotation downloaded from Ensembl. Sleuth (v0.30.0) was used for between sample normalization and to calculate aggregate gene counts.

### Gene expression analysis using QuantSeq

Peripheral blood cells were isolated from WT and miR-144^Δ/Δ^ zebrafish 2-dpf and adult zebrafish in biological triplicates. Total RNA was extracted with TRizol and quantified. Libraries for sequencing were prepared using QuantSeq 3’ mRNA-Seq Library Prep Kit FWD. QuantSeq libraries were analysed as described above.

### Single-cell RNA sequencing of the hematopoietic niche of Danio rerio whole-kidney marrow and dorsal aorta

Pronephros of adult zebrafish was isolated according to previously described protocol^63^. Fish to be dissected were approximately age-matched and only males were used for each condition. At the time of dissection, fish was euthanized in ice-cold water for two minutes until all breath ceased and its head was removed by making a cut immediately behind the gill operculum using a surgical blade. It was placed on a slightly wet sponge with a notch to maintain the fish on its back. Using a surgical blade, a ventral midline incision was made from anterior to posterior to expose the body cavity, which was held open with forceps. With a second pair of forceps, the body cavity was removed of any sperm, digestive organs, or other tissue. Upon successful cleaning of the body cavity, only the whole kidney marrow (WKM) and dorsal aorta were visible attached to the dorsal body wall of the opened fish. The WKM and attached dorsal aorta were carefully removed using forceps and added to a microcentrifuge tube with 1mL ice-cold 0.9 X PBS containing 2% FBS and 5mM EDTA, which was then maintained on ice.

The sample was resuspended by vortex as pipetting up and down. The entire sample was pipetted onto a 40 micron mesh (Falcon) over a 50mL Falcon tube and crushed with the plunger of a 1mL syringe. Cells were pelleted by centrifugation at 200 x g for 5 min at room temperature. Cells were resuspended in 1mL of ice-cold 0.9 X PBS containing 2% FBS and 5mM EDTA, and transferred to a 1.5 mL microcentrifuge tube. The cells were pelleted once again by centrifugation at 200 x g for 5 minutes. Upon aspiration of the supernatant, the pellet was resuspended in 500 µL of ice-cold 0.9 X PBS containing 2% FBS and 5 mM EDTA and passed through a 35 micron mesh cell strainer cap into a 5 mL polystyrene round-bottom tube (Falcon). To the resuspension, 1 µL of 1 mM TO-PRO-3 Iodide in DMSO (Thermo) was added to stain dead cells and 1uL of DRAQ5 (Invitrogen) was added to stain viable cells. A Beckman Coulter MoFlo Astrios cell sorter was used to gate-select for viable cells of the hematopoietic niche, while excluding non-hematopoietic kidney cells. These cells were sorted out into 1.5 mL microcentrifuge tube containing 50 µL of ice-cold 0.9 X PBS containing 2% FBS and 5mM EDTA.

The sorted samples were kept on ice and a fraction was used to count the cells in a hemocytometer. The sample was then processed for single-cell RNA sequencing. Single-cell sequencing was performed using 10x Genomics platform at Boston University Microarray and Sequencing core.

scRNA-Seq data was analyzed as follows: Reads were aligned to the zebrafish genome (GRCz11) with STAR (v2.7.9). Ensembl (v104) gene annotations were downloaded as a GTF file and used for the analysis. Results files in h5ad format were read into Python and labelled as WT or miR-144 and then combined into a single Anndata object. Scanpy was used for normalization and filtering. After creating the UMAP and clustering it was observed that one outlier cluster was enriched for genes expressed in sperm and was removed as it was likely contamination on sample collection. Clusters were annotated with the set of marker genes used in ^64^ and progenitor cell and erythrocyte clusters were further analyzed to compare the expression of hmgn2 in the wildtype and miR-144 mutant genotypes.

### Cloning, Lentivirus Production and Infection

We designed an Hmgn2-3’UTR reporter based on the pHAGE lentiviral vector that has been previously described^65,66^ DNA sequences encoding human Hmgn2 coding region, EYFP and either wild-type (*hs*Hmgn2-EYFP-3’UTR-WT) or mutant lacking miR-144 target sites (*hs*Hmgn2-EYFP-3’UTR-MUT) 3’UTRs were ordered directly as gBlocks (IDT) and inserted into pHAGE vector at BamH1 and Not1 sites. Lentiviruses were produced using a five-plasmid transfection system in 293T packaging cells as previously described^65^. Supernatants were collected every 12 hours on two consecutive days starting 48 hours after transfection, and viral particles were concentrated by centrifugation at 16,500 rpm for 1.5 hours at 4°C. Immortalized iPSC-derived erythroid cells were infected with 15 μl of concentrated virus in the presence of polybrene (5 μg/ml). The medium was replaced after 16 hours with base Serum-Free Expansion Medium I (SSI) supplemented with 2 mM of L-glutamine, and 100 μg/mL of primocin 40 ng/mL of IGF1, 5 × 10^−7^ M of dexamethasone, and 0.5 U/mL of hEPO.

### Erythroid differentiation of human iPSCs

Cells were derived from a library of human sickle cell disease induced pluripotent stem cells (iPSC)^67^. Differentiation of these iPSCs faithfully recapitulate human erythropoiesis in vitro^68^. Hematopoietic differentiation from iPSCs to hematopoietic stem and progenitor cells (HSPCs) was induced according to Vanuytsel *et al*.^35^. HSPCs were then transferred into SSI medium (see below) to induce erythroid specification. After 5 days into SSI medium, when the cells reached a stable proerythroblast stage, a lentiviral vector carrying large T antigen was introduced to immortalize cells. Immortalized cells were subsequently infected with lentivirus encoding *hs*Hmgn2-EYFP-3’UTR-WT, *hs*Hmgn2-EYFP-3’UTR-MUT or EYFP alone. To induce further erythroid differentiation from what we consider baseline (’day 0’), cells were cultured for 2 more days in SSI media (‘day 2’), followed by 5 days into SSII media (see below, ‘day 7’). SSI and SSII media are parts of a 2-step suspension culture system with a base medium consisting of StemSpan Serum-Free Expansion Medium II (StemCell technologies), 2 mM of L-glutamine, and 100 μg/mL of primocin. To create SSI medium, this base was supplemented with 100 ng/mL of human stem cell factor, 40 ng/mL of IGF1, 5 × 10^−7^ M of dexamethasone, and 0.5 U/mL of hEPO and cells were cultured at 37°C in normoxic, 5% carbon dioxide conditions. SSII medium is created by adding 4 U/mL of hEPO to the base medium. Cells were cultured in hypoxic conditions during the maturation phase in SSII.

### cDNA Preparation and qPCR

RNA was extracted using the Trizol (Qiagen) and DNase treated using a DNA-free kit (Ambion). Complementary DNA (cDNA) was prepared from total RNA (5 μg) by reverse transcription using LunaScript^®^ RT SuperMix Kit (NEB). qPCR reactions were performed using Power SYBR Green Master Mix (Thermo) and carried in ViiA7 Real-Time PCR System (Applied Biosystems). Data were normalized to *GAPDH* mRNA amplification. For the expression analysis of globins in human cells predesigned TaqMan primers (Applied Biosystems) were used in conjunction with Taqman Universal Master Mix II (Applied Biosystems; #4440038) for quantitative reverse transcription PCR analysis on the StepOne/QuantStudio 6 Flex Real Time PCR Systems (Applied Biosystems). The housekeeping gene *GAPDH* was used as an endogenous control. Data were analyzed using the ddCT method. Fold expression was calculated as 2^−(ΔCt)^, with ΔCt = Ct(gene of interest) − Ct(β-actin). These normalized fold expression values for all time points were then divided by the day-0 values to obtain relative fold expression compared with day 0 of erythroid differentiation.

### Flow cytometry analysis

Cells were stained on ice for 25 minutes using the following antibodies: CD235a-PE (BD; #555570), Flow cytometry was conducted on a Stratedigm S1000EXI. FlowJo v8.7 (FlowJo, LLC) software was used for analysis, and FACS plots shown represent live erythroid cells based on side-scatter/forward-scatter gating.

### Small RNA Northern blotting

Total RNA was extracted using Trizol (Invitrogen), quantified (we used 10 μg of total RNA per lane) and resuspended in formamide. Loading buffer 2X (8 M urea, 50 mM EDTA, 0.2 mg/ml bromophenol blue, 0.2 mg/ml xylene cyanol) was added and the samples were boiled for 5 min at 95°C. miRNAs were separated in 15% denaturing urea polyacrylamide gel in 1X TBE and then were transferred to a Zeta-Probe blotting membrane (Bio-Rad) using a semi-dry Trans-Blot SD (Bio-Rad) at 20 V (0.68A) for 35 min. Membranes were UV cross-linked and pre-hybridized with ExpressHyb Hybridization Solution (Clontech) for 1 h at 50°C. Membranes were blotted with 5′ ^32^P-radiolabelled DNA oligonucleotide probes at 30°C overnight. Membranes hybridized with oligonucleotide DNA probes where washed at room temperature with 2X SSC/0.1% SDS followed by 1X SSC/0.1% SDS for 15 minutes. The blots were exposed to a phosphorimaging screen for 1 to 3 days. Signal was detected using the Typhoon IP phosphorimager (GE Healthcare Life technologies) and analyzed using the ImageQuant TL software (GE Healthcare).

### Preparation of radiolabeled probes

Radiolabeled DNA probes were prepared according to the StarFire method. Oligos carrying specific DNA sequence complementary to the miRNA of interest were annealed to the universal oligo (5′-TTTTTTTTTT666G6(ddC)-3′, where “6” corresponds to a propyne dC modification) via complementary hexamer sequence. Annealed duplexes are then labeled with α-^32^P-dATP (6 μL of 10mCi/mL stock) using the Klenow fragment of DNA polymerase. Reaction was stopped by adding 40 μL of 10 mM EDTA solution to 10 μL of reaction. Then labeled oligos were purified using Micro-Spin G25 columns (GE HealthCare). 3,000,000 cpm of the P^32^ labeled StarFire probes were used to probe the membrane.

### Western blotting

Proteins were resolved using precast NuPAGE Novex 8% Bis-Tris gel plus (Invitrogen) at room temperature and constant voltage (100 V) in 1X MOPS Running buffer and transferred to a nitrocellulose membrane (0.45 mM, Bio-Rad) using iBlot2 system (Invitrogen). After transfer membranes were blocked in a blocking buffer (5% non-fat dry milk in TBS-T) for 1 hour at room temperature. Membranes were incubated with primary antibody overnight at 4C with following dilutions. Anti-HA (1:1000) anti-actin (1:1000). After that membranes were washed three times with TBS-T and then incubated with secondary antibody for 2h at room temperature with following dilutions. IRDye 800CW Goat anti-mouse (1:10000), IRDye 680RD Goat anti-rabbit (1:10000). All membranes were scanned using Odyssey Scanner (Li-COR) and band intensity was quantified with Odyssey software.

---

**Figure S1.**
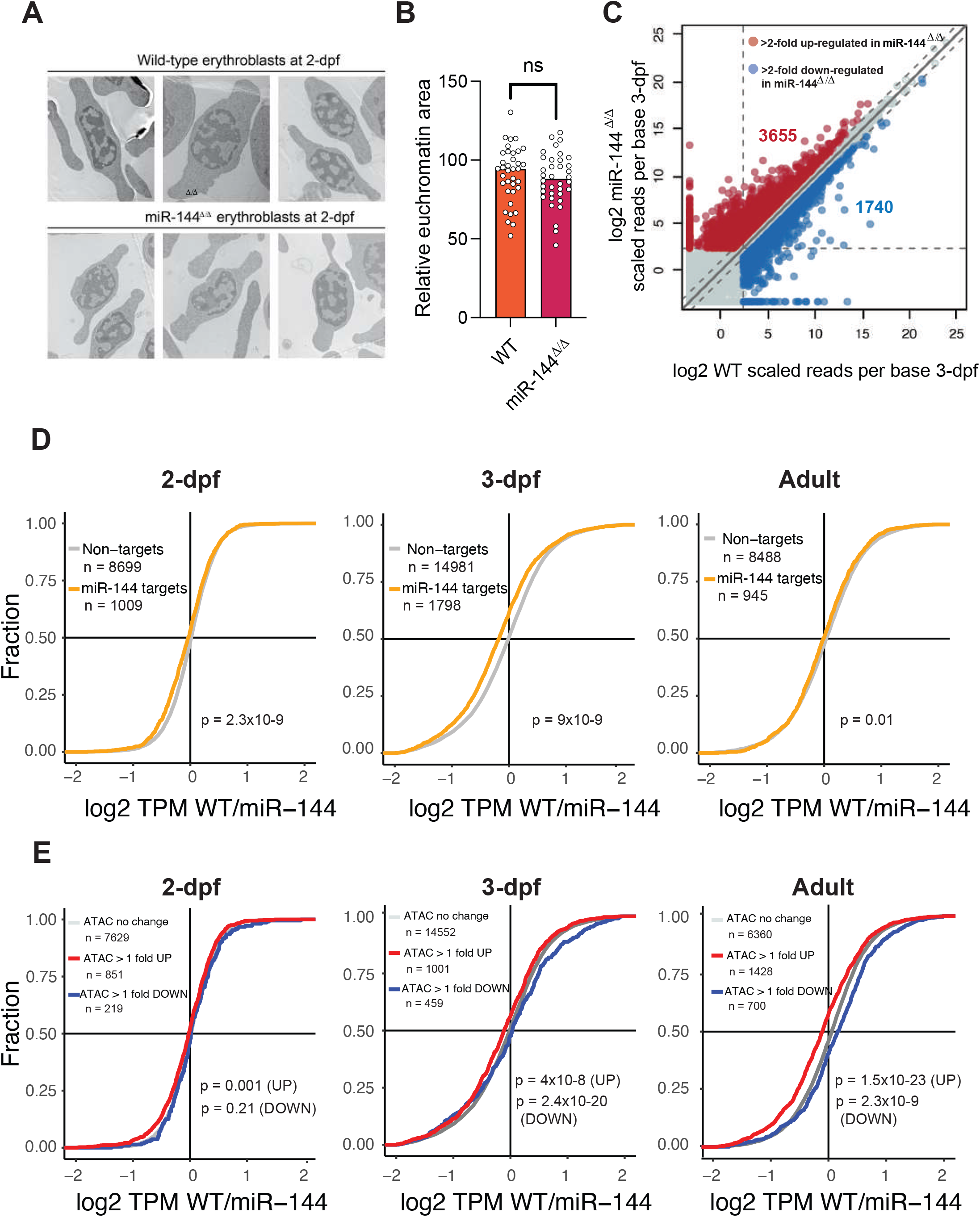
**(A)** Transmission Electron Microscopy of erythrocytes isolated from 2-dpf embryos. **(B)** Quantification of euchromatic regions of the nuclei from (A). 35 cells are analyzed in each case. *p*-values from unpaired *t-*test. Error bars represent standard error of the mean. **(C)** RNA-Sequencing of erythrocytes isolated from 3-dpf from miR-144^Δ/Δ^ zebrafish and wild-type siblings. Average of 2 biological replicates is plotted. **(D)** Cumulative distributions of fold changes revealed by RNA-Sequencing between wild-type and miR-144^Δ/Δ^ and wild-type erythrocytes for curated miR-144 targets (expressed more than 5 scaled reads per base in erythrocytes and have predicted miR-144 targets sites according to TargetScan (red line) and non-targets (grey). **(E)** Cumulative distributions of fold changes of gene expression revealed by RNA-Sequencing between miR-144^Δ/Δ^ and wild-type erythrocytes for the genes with increased more than 1-fold (red line), decreased more than 1-fold (blue line) and not changed (gray line) chromatin accessibility in genes.

**Figure S2.**
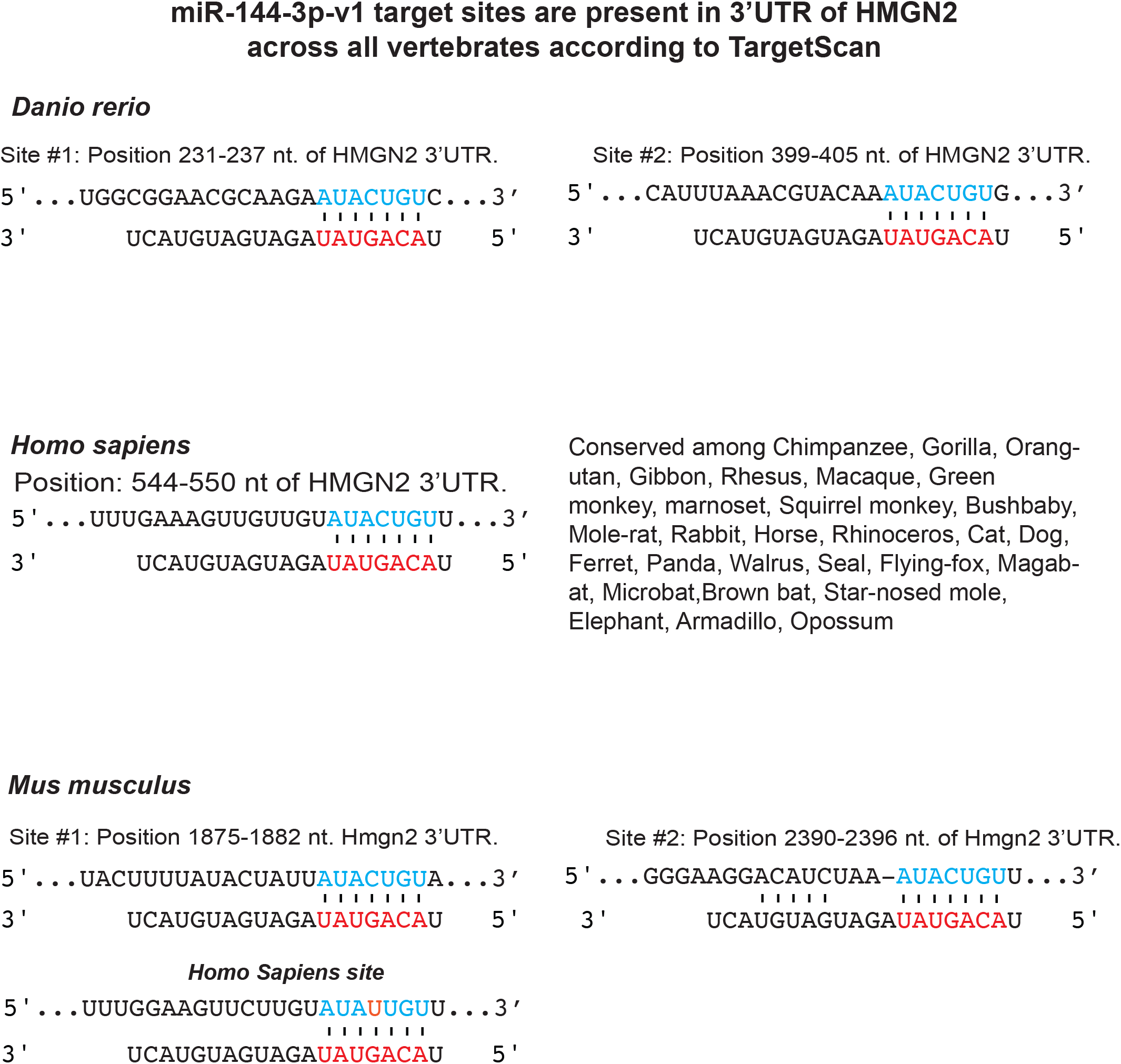
Predicted miR-144-3p-v1 (according to TargetScan target sites in *Hmgn2* 3’UTR of *Danio rerio*, *Homo Sapiens* and *Mus Musculus*. miR-144 seed region is indicated in red.

**Figure S3.**
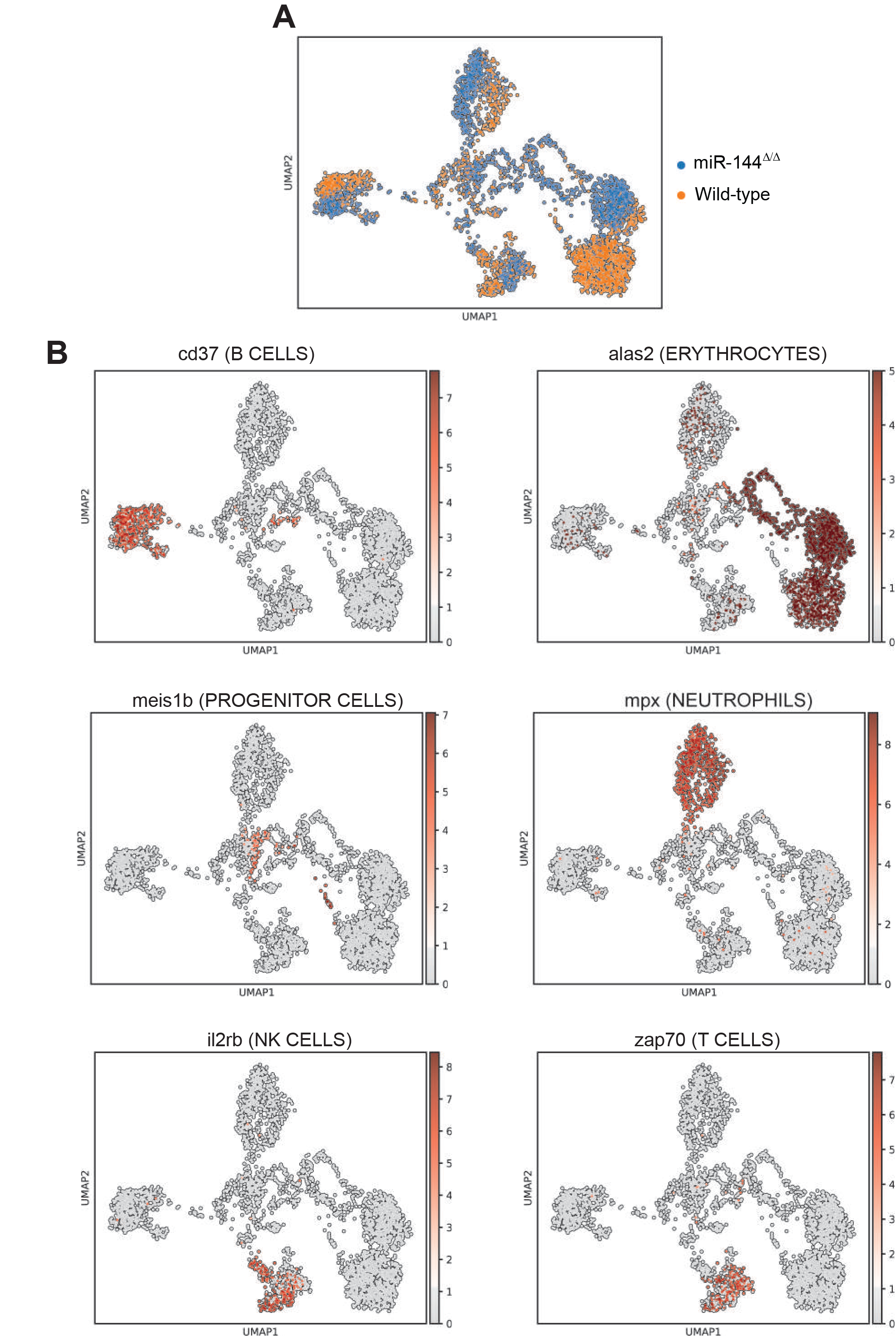
UMAP of scRNA-Seq data from zebrafish pronephros. **A)** UMAP colored by genotype, each dot represents a single cell. **B)** UMAP colored according to the expression of marker genes representative of each main hematopoietic cell lineage.

**Figure S4.**
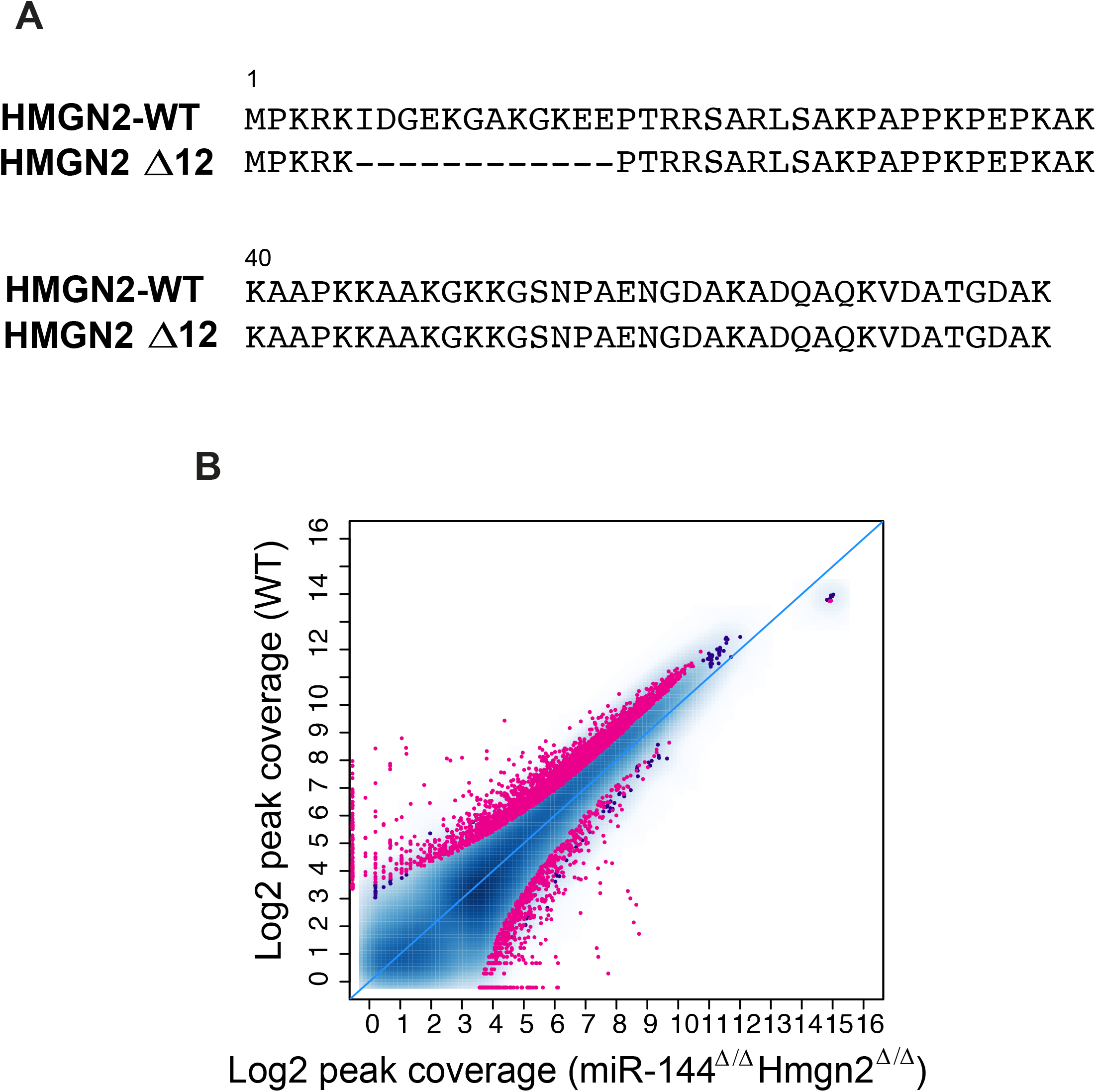
**A)** Protein sequence alignment of wild-type and mutant zebrafish Hmgn2. **B)** Scatter plot showing differential chromatin accessibility in miR-144^Δ/Δ^ and miR-144^Δ/Δ^ Hmgn2^Δ/Δ^ using ATAC-Seq. Each dot represents an ATAC-seq peak.

